# Characterising Protein Search Drift using exhaustive protein search and Alphafold2

**DOI:** 10.1101/2024.11.14.623594

**Authors:** Daniel WA Buchan

**Affiliations:** UCL

## Abstract

In this paper we present the first exhaustive analysis of iterative protein search drift and show how such results may impact downstream modelling. Assembling and extracting evolutionary information from families of related proteins is a core challenge in the studey of molecular evolution. For instance, iterative protein search is a common first step in a wide variety of bioinformatics tools and pipelines. And the output of such searches often form the inputs for modelling tools such as Alphafold2. Here we characterise profile drift; the tendency for some searches to become contaminated with sequences outside of the intended evolutionary family. We observe that drift occurs in nearly 15% of searches and can be observed to have measurable impacts on downstream predictive tasks such as structure prediction.

## Introduction

The motivation for this work is to attempt to understand some classes of unwanted or non-optimal behaviours during protein database search. Established methods of protein database search include a number of iterative and non-iterative methods such as FASTA(1), BLAST(2), CLUSTAL-Omega(3), PSI-BLAST(4), HH-SUITE(5) and HMMER(6). With such tools, a query sequence is searched against a database of entities which are designed to encode a large database of target proteins. Such entities can either be single protein sequences (e.g FASTA), a compressed search structure representing that database (e.g. BLAST/PSI-BLAST) or set of models which represent the protein database (e.g. HH-SUITE). The goal of protein sequence search is typically to identify the subsets of proteins within the database which are evolutionarily related to the query sequence. After an initial search hits can be accepted as evolutionarily related if they pass some scoring or statistical threshold set by the user. In a non-iterative algorithm such as FASTA this set is the complete output of the search. In iterative search methods (*i*.*e*. PSI-BLAST), the initial hits can be regarded as forming a putative evolutionary family with the query sequence and this family can be used for further iterations of the search procedure. The set of proteins are converted into a model or profile of the evolutionary-family which represents the sequence diversity present in the sequence family members. These models can then be used to search the database a subsequent times.

In the case of PSI-BLAST the intermediary model that is compiled at each iteration is a called a Position Specific Score Matrix or PSSM. In the case of HHSuite and HMMER a Hidden Markov Model (HMM) is produced to represent the family of sequences. If we consider that the growing family of proteins can be converted to a multiple sequence alignment (MSA), then the goal of building theses intermediary models is to represent the statistical variation in the amino acid (or nucleotides) in each column of the MSA. In the case of the PSSM the matrix columns have one-to-one concordance with the columns in such an MSA.

For iterative search methods these models are used as the basis for subsequent iterations of the database search. The logic here is that conditioning the search using the statistical variation contained in the family will be better able to find more distant evolutionary relatives in the subsequent iterations. Iterations can be completed an arbitrary number of times to search for successively more distant proteins. But typically, search is carried out for some fixed number of iterations or until the results converge where no new protein sequence hits are identified. Once all iterations are complete a final such model can be built and stored as a statistical representation of a “complete” evolutionarily related protein family. There models can be thought of as a “Sequence Family Profile”. Databases such as Pfam (1)make these models available to users.

Ideally, a protein family is the representation of a cluster of proteins that forms a discrete group within the space of known protein sequences. Often the goal of protein search is to discover a set of evolutionarily related proteins “near” the query sequence such that they share some important, salient biological property such as shared biochemical function, 3D protein structure or a common evolutionary ancestor. However, it is observed that membership of a search cluster changes as the iterative search progresses. Usually this is the desired behaviour of the search process. That is, to expand the set of discovered proteins to find more distantly related proteins. However, some changes are unwanted such as, including proteins that do not appear to be in the same evolutionary family, are not functionally related or do not share the same structure. If we imagine that search produces a cluster of related proteins then including members which are too distant can be imagined to be moving the search cluster centroid away from the intended regions of sequence-space (*i*.*e* away from the query sequence). Such non-optimal behaviours have been previously identified as “profile drift” (7,8). That is, the protein family model that is constructed at each iteration is drifting away from either the query sequence or away from the putative target region of protein sequence space (see Figure 1).

**Figure 1:**
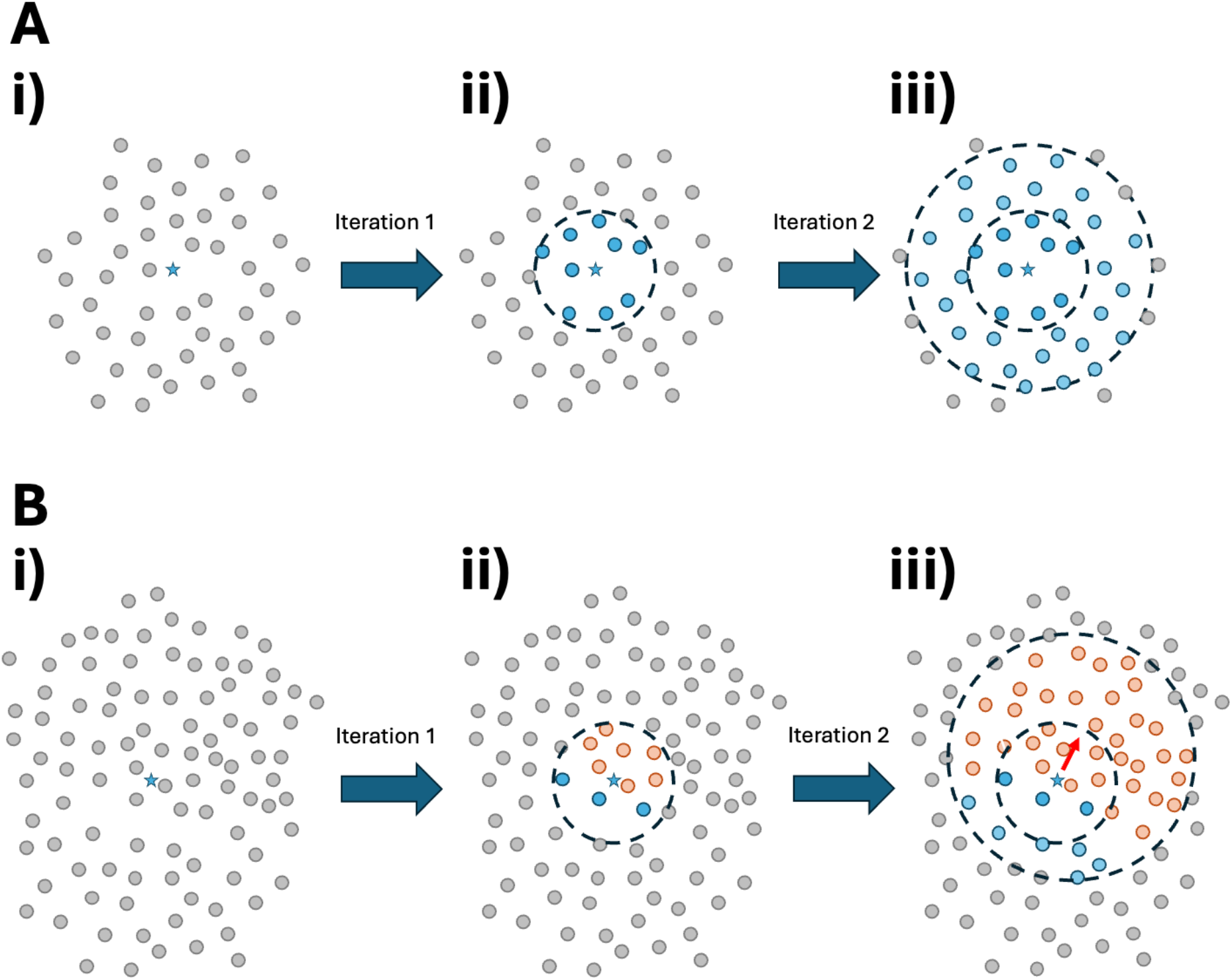
Two examples of protein search in a figurative protein sequence space. Protein sequences are visualised as grey spots in a 2D space such that their evolutionary distance is analogous to the Euclidean distance in this projection. In **A** we visualise an “expected” behaviour for protein search. We start with some query sequence (the blue star) located in the sequence space. In Iteration 1 (moving from **Ai** to **Aii**) we identify evolutionarily “nearby” sequences which are all of the same sequence family (blue) as the query sequence, as desired. These can then be constructed into a model of the sequence family that is used in Iteration 2 (**Aii** to **Aiii**) to find more distantly related sequences from the same family. Iterative search drift is illustrated in **B**. Here the sequence space is more densely populated in one direction. On the first iteration of search sequences nearby the query sequence (blue star) are found. Here the hits are from two different sequence families, blue and orange. The orange sequences out number the blue target family sequences so the profile that is built for iteration 2 is dominated by the sequence diversity from the orange family. In **Biii** we see that this may cause more orange family sequences to be recruited thus also moving the centroid of this sequence space cluster away from the query sequence (red arrow).

This profile drift or profile contamination effects can have impacts across many forms of bioinformatics analyses. The ability to discover coherent sets of sequence relatives is a critical part of many predictive methodologies in bioinformatics. These include protein function prediction methods such as PFP(9), INGA(10) and FFPred(11), protein classification methodologies such as CATH(12), PFAM(13), and proteins structure prediction algorithms such as AlphaFold2(14), DMPfold2 (15)and OpenFold(16). When trying to make accurate predictions using protein families including “off-target” family members may increase the number of false-positive predictions in classification tasks or increases error rates in modelling tasks.

However, defining drift runs into important philosophical questions about the nature of protein families. Firstly, protein space is not evenly populated, some protein families are more populated (*i*.*e*. are more exhaustively explored by evolution) and have greater sequence diversity than others. Take for instance the Rossmann fold protein domain, this is one of the most commonly replicated proteins domains known (17,18) and such domains are common found in enzymes. Immunoglobulin fold domains are ubiquitous in cell-signalling proteins. There is an open question to what extent such densely populated regions of sequence space should be divided in to separate families. And what functional, structural and evolutionary criteria should be used to delineate where to draw the family boundaries. Designations such as Pfam Clans (19) seek to acknowledge the complex inter-relationship of sequence families in dense regions of protein sequence space.

Another feature of protein families is that they are not discretely spaced apart in “protein space”. Some sequence families are obviously have large evolutionary distances separating them. On the other hand, P-loop NTPase domains share evolutionarily heritage with other Rossmann Fold families (see CATH fold 3.40.50 and PFAM CLAN CL0023) when families are evolutionarily close this introduces a degree of ambiguity where one sequence family begins and the next ends. Where should we draw the line about which sequence belongs to which domain family? How many homologous families should the Rossman folds be divided in to?

Additionally, as with all biological classification, a protein family is a human construct and the criteria we use to define a family may not be the best way to represent the evolutionary processes that gave rise to differing sequences. For instance, it is certainly scientifically useful that the CATH database segregates it’s Homologous Protein Families into functionally homogenous groups but the evolutionary trajectory of the proteins was not opimised to make tidy, discrete families. From a purely evolutionary perspective it may not make sense to divide some closely related sequences in to a separate families, as functional changes can be the consequence of very few mutations. But human classification decisions can be guided by their utility too, such as aiding us when attempting to build systems for protein function prediction.

An upshot of these considerations is that any analysis of protein search drift will necessarily pick up artefacts of the classification methodology as well as the drift issues that arise from the statistical errors inherent in the search methodology. That is, from a statistical and evolutionary point of view, it may be completely valid that a search for P-loop NTPAses pulls in many sequence-related Rossman fold domains in it’s results. Even though the person performing the search may only wish to find other P-loop NTPases. But it is unlikely we can trivially disentangle which types of drift we see are statistically incorrect and which are a function of the real density of the region of protein space we are searching. Using structure prediction as a “gold standard” we attempt to estimate this in the following paper.

In the first part of this study we look to characterise protein drift. Using the commonly understood tool PSI-Blast, we conduct a large number of experiments to measure how often iterative search includes out-of-target-family sequences to the search results. This allows to explore how often search drift happens and and if there are different classes of drift behaviour. This also gives us a view on how often a PSI-BLAST PSSM may represent a discrete, region of the search space. We then examine how drift impacts downstream predictive tasks. Alphafold2 is a common and important, contemporary predictive algorithm. We show how the reliability of the models produced is impacted for each category of drift across the iterations of the search results. Finally, we investigate how coherent, in comparision to PSI-BLAST PSSMs, the representation of protein families is for some other types of sequence models such as Hidden Markov Models (HMMs) and Large Language Models (LLMs).

## Method

In the first phase of the study we look at how iterative protein search typically progresses. PSI-BLAST was chosen as it is a commonly available tool for searching protein datasets. It is widely used, understandable and it is trivially easy to monitor the outputs and behaviour of its iterative search process. The fasta dataset for version 100 of the Pfam database was downloaded. This contains sequence data for 20,049 Pfam domain families. For each Pfam family a single sequence was randomly selected to act as representative sequence (the Rep) for that family. To investigate drift each Rep was then searched against the whole Pfam v100 fasta sequence database using PSI-BLAST. PSI-BLAST was run with default parameters and searched for 20 iterations. 20 iterations is beyond what is typically needed but this was selected to ensure all searches had a chance to either converge or saturate. At each iteration we capture the set of hits that were identified at that iteration and record which Pfam family each hit belongs to.

The aim here is to compile a list of families where we witnessed profile drift. That is, where the hits from each iteration of the search include sequences which do not share the same Pfam family membership as the Rep which was used to initiate the search. 5,776 families were seen to have drift and this comprises just less than 30% of all searches. The trajectories of the searches were analysed and this indicates that search trajectories cluster in to roughly 6 groups which display differing kinds of drift behaviour. This is discussed in full in the Results below. For families that show drift behaviours the amount of out-of-family sequence gives a measure of how “coherent” the final PSSM model for that family is.

To investigate the density of sequence space the Pfam Reps were used to perform an all-against-all comparison alignment between all 20,049 Reps using the EMBOSS 6(20) implementation of Needleman and Wunsch(21). Every alignment score was recorded and added to 20,049 × 20,049 similarity matrix. This matrix was normalised to between 0 and 1 and it was converted to a distance matrix by taken 1-*x* for each cell in the matrix

Next, to observe how the search results for each class of drift impacted a downstream prediction task up to 100 Pfam families from each drift class were randomly selected. The previously calculated sequence hits for each PSI-BLAST iteration for each family search were retrieved. It is possible for a family to have fall in to two classes of drift and some classes of drift affected fewer than 100 pfam families. This resulted in 530 Pfam families being selected. Clustal Omega, was used to generate an MSA for the hits for each iteration for each family. Clustal Omega was run with its default settings. Using CollabFold(22) we generated 3 Alphafold2 models using these MSAs. CollabFold was adapted to run Alphafold2 only with MSA input and with no template search. For every Pfam family in each drift class, one PDB model was built for the iteration 1 hits and one at iteration 20. For each of the 4 drift classes where out-of-family contamination is observed a 3^rd^ model was generated for the iteration with the greatest percentage of out-of-family hits. For the 2 drift classes that showed limited contamination a 3^rd^ model was generated for iteration 5 (see discussion in Results below). This results in 3 PDB models for each Pfam family for each drift class. Change in mean plDDT for each model was calculated.

To test if changes in mean plDDT were also associated with in changes in structural family or protein fold merizo-search was run for each every Alphafold2 model. The CATH Homologous superfamily the model belonged to and the max TM-Score between the model and closest CATH domain match was recorded. As each model is a single Pfam domain merizo-search was used in search mode with standard parameters.

The final test seeks to compare how “coherent” other easily available forms of protein family representation are compared to the PSI-BLAST PSSM. The PFAM database provides a full suite of HMM profiles for each of it’s protein sequence domain families. Though the seed alignments are provided they do not provide information sequences were added to the HMM at each stage of the iterative HMMER search that was used to prepare each final HMM. Seed alignments used to prepare the whole sequence family are, in-part, built with manual curation. So the process to build these profiles is not quite analagous to a simple iterative search process like PSI-BLAST. Nevertheless the resulting library of HMMs are pre-calculated models of sequence diversity on a per-Pfam family basis. HMMER’s hmm-emit tool was used to generate new “in-family” sequences. From there we can measure whether or not these putative new sequences do belong to the family the HMM Profile represents. To this end the hmm-emit was used with previously selected 530 families to generate 50 new sequences for each family.

To test if these new sequences did belong to the family in question FASTA was used to match the new sequence against the Pfam v100 fasta sequence database. For each FASTA search the Pfam family membership of the top hit was taken as the most likely family identified for the newly generated sequence. This was used to as a measure of whether new synthetic sequence was indeed in-family.

Finally, sequence generation process was repeated for the the T5ProtTrans protein Large Language model, pLLM(23). The masked-sequence training task used to prepare the pLLM should embed information about sequence variation within sequence families in the weights of the model. The ProtTransT5 model was downloaded and used it to generate new sequences for the 530 random PFAM drift families selected above. To generate a new sequence for a family we randomly select a sequence from one of the pfam families and mask a given percentage of its residues. This masked sequence is used as input to the network and the inference step is run to generate an “unmasked” sequence. This is essentially equivalent to the training task that was used to train the model. Our objective is to discover to what extent the model can reconstruct a valid in-family sequence. 50 new sequences were generated for each family. And this process was completed 3 times with different levels of residue masking; 25, 50 or 75 percent

## Results

Table 1 shows the headline results for the 20,049 PSI-BLAST runs. A majority, 71%, of PSI-Blast searches only find hit sequences, across all 20 iterations of search, from within the same Pfam family as the Rep which was used to initiate the search. This indicates that drift behaviours, where sequence searches find sequence relatives that are outside their pre-assigned sequence family, occur in only 29% of searches.

**Table 1:** Drift statistics showing the number of Rep PSI-BLAST runs calculated and the breakdown of how many PSI-BLAST runs included hits that were outside the Rep’s Pfam family (drift behaviour) and how many runs only included hits that were in the Rep’s Pfam family (no drift behaviour).

|  | Count |
| --- | --- |
| Rep PSI-BLAST Runs | 20,049 |
| Results with no drift behaviour | 14,273 |
| Results with drift behaviour | 5,776 |

These results were analysed to observe how drift occurs during these searches. 6 broad classes of drift behaviours were observed. Firstly “No Drift, where the search process only recruits sequences that belong to the same sequence family as the Rep query sequence. “Minimal Drift”, where no more than 5% of the hits are from different sequence families as the query Rep. “Query Purified” where the hits from the same sequence family that the query Rep belongs to are eliminated from the search results as the iterations progress. “Contaminants Grew”, where hit from sequence families other than query Rep family are observed and their proportion of the results increases as then search iterations continue. “Contaminants Purified” where the hits from out-of-family sequences appear and make up more than 5% of hits but search iterations increase they are eliminated from the results. And lastly “Contaminants Complex” where there are out-of-family hits across many contaminant sequence families but they follow more than one of the above contaminant behaviours observed. This is summarised in Figure 2.

**Figure 2:**
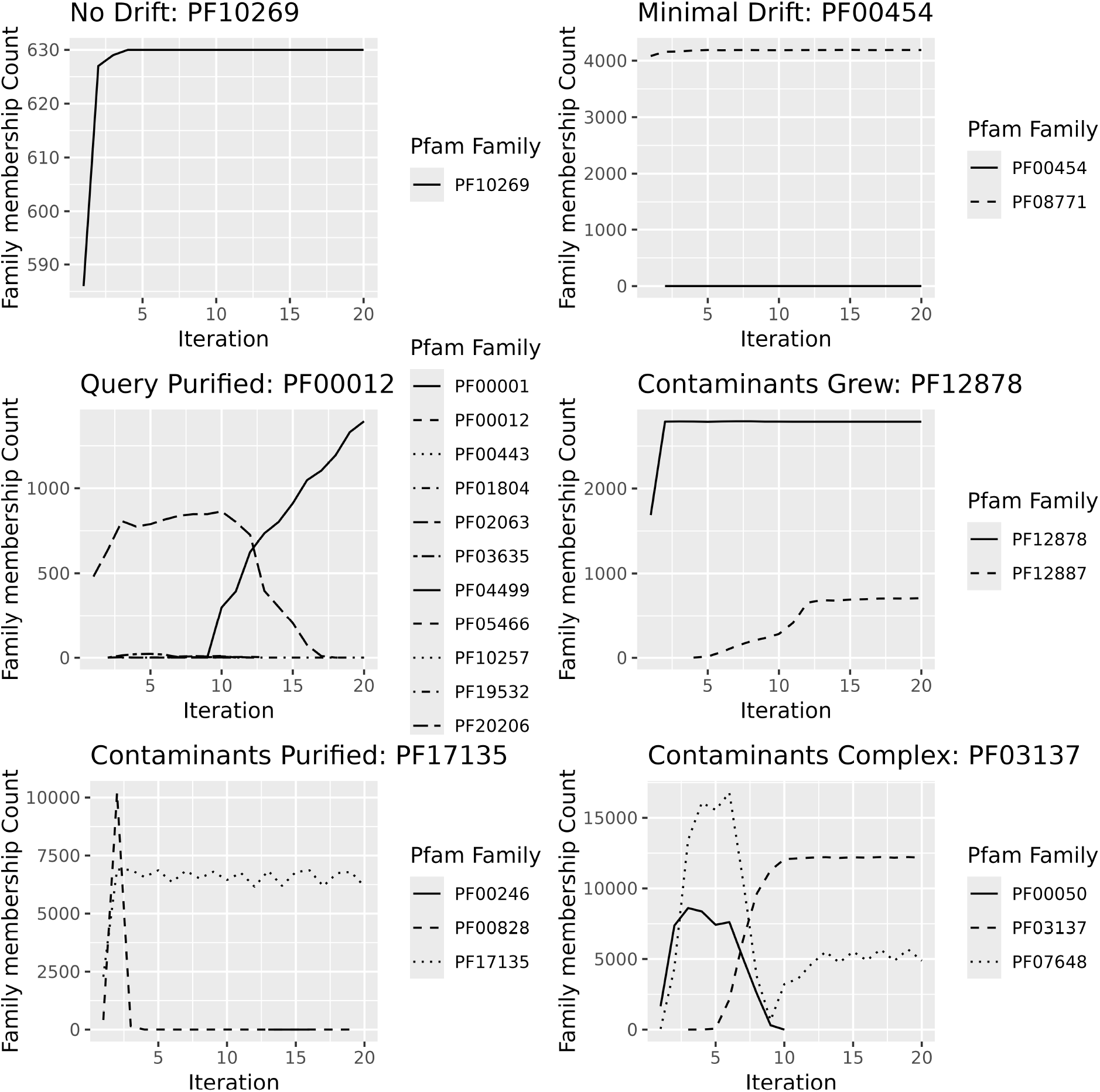
Examples of each class of drift which was identified with the iterative PSI-BLAST search. There are 6 classes “No Drift”, “Minimal Drift”, “Query Purified”, “Contaminants Grew”, “Contaminants Purified”, “Contaminants Complex”. Description of these classes can be found in the text.

Having identified these search/drift behaviours each Pfam search can be classified into one of the categories. It is worth noting that each search can fall in one or more of these categories and they are not mutually exclusive. Table 2, indicates a summary of the search categories. One important result here is that nearly 5% of searches resulted in the sequence family that the query belongs to being eliminated from the search results. That is, in terms of a sequence family cluster, the centroid of sequences cluster has moved away from the Rep sequence family to some other region of sequence space. In nearly 15% of the searches once a significant number of off-target sequences appear in the results they remain a feature of the results and typically grow as a percentage of the search results. Again indicating that the centroid of the search cluster has moved away from the target family. In total the minimal drift, non-drift groups make up 17,021 of the search cases indicating. around 85% of searches proceed with minimal of contamination.

**Table 2:** Counts of the searches by drift category.

| **Drift Class** | **Count** |
| --- | --- |
| Non-Drift | 14,273 |
| Minimal Drift | 2,748 |
| Query Purified | 1,063 |
| Contaminants Grew | 2,044 |
| Contaminants Purified | 50 |
| Contaminants Complex | 924 |

Analysing the results by iteration what is seen is that the category of drift a search falls in to is set by the 10^th^ iteration of search and in most cases before the 6th. That is, when either the contaminant families or Rep family are eliminated from the results it is commonly prior to iteration 10 in the results. Or where the number of contaminants is large or complex this is seen in the results prior to the 4^th^ iteration. One observation here is that there is limited utility in running iterative search beyond 5 iterations and in about 5% of cases sequence from the target family will be fully eliminated by that time (this information is summarised in see table 3). When non-Rep family sequences are recruited to the search results, the mean number of off-target families seen is 4.75. Which could indicate a dense region of sequence space with many neighbouring or overlapping families.

**Table 3:** mean iterations during search that various contamination “events” are observed.

| **Class** | **Mean Iteration Observed** |
| --- | --- |
| Query Purified from search results | 5.97 |
| Contaminant family purified search results | 9.8 |
| Contamination of more than 5% (Grew and Complex) | 3.9 |

It is worth considering if drift behaviours are the consequence of the the density of the sequence space being search. We can examine the distance matrix that was prepared to answer this question. The modal average distance between Pfam families in the distance matrix is 0.97 with a standard deviation of 0.25 indicating that there are typically only large distances (in terms of relative Needleman and Wunsch scores) between any random pair of pfam family reps. Examining the pfam families where drift is observed the average distance from the query family to any contamination family is 0.91, nearly 2 standard deviations away from the mode. This suggest that contaminant families are closer to the query sequence than the typical pfam family. Somewhat supporting the notion that drift behaviours are a consequence of more densely regions of protein sequence space.

The next experiment seeks to characterise the extent to which each drift category impacts downstream predictive tasks which rely on assembling accurate, discrete families of protein sequences. 100 families were sampled from each drift category (only 50 for the contaminants purified class), the sequences were aligned built Alphafold2 models were built with the alignments. Table 4 summarises the average plDDT scores across the selected set of iterations. Here was see that typically Alphafold2 is robust to the changes in alignment composition. The Non-drift and minimal drift categories see only a slight increase in the calculated plDDT over 20 iterations of search. This is likely explained by the fact that on average the MSAs contain a greater number of effective sequences at iteration 20 than at iteration 1. The most significant impact of drift is in the Query Purified class, a 10 point drop in mean plDDT is observed in this instance. This indicates the changing composition of the MSA has a large effect on the reliability of the structural model in these cases. The behaviour of those drift classes which contain both in-family and contaminating family hits appears are more complex to interpret.

**Table 4:**
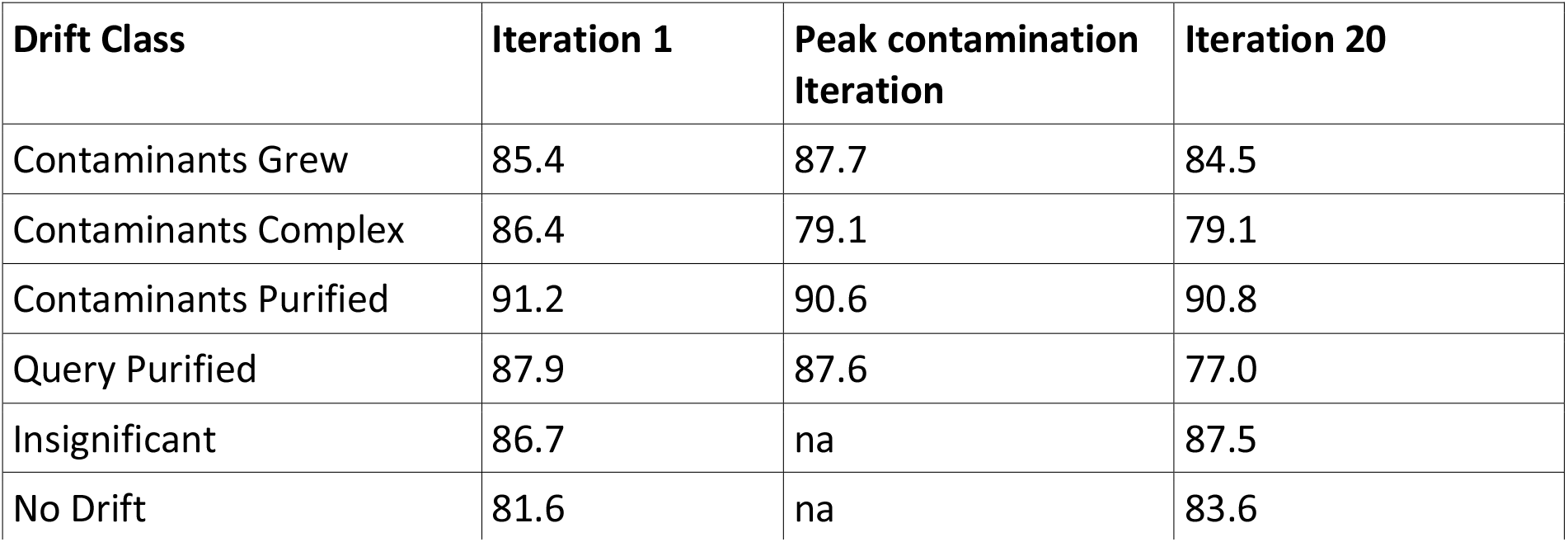
summary of mean plDDT for AlphaFold2 models created for each drift class.

| Drift Class | Iteration 1 | Peak contamination Iteration | Iteration 20 |
| --- | --- | --- | --- |
| Contaminants Grew | 85.4 | 87.7 | 84.5 |
| Contaminants Complex | 86.4 | 79.1 | 79.1 |
| Contaminants Purified | 91.2 | 90.6 | 90.8 |
| Query Purified | 87.9 | 87.6 | 77.0 |
| Insignificant | 86.7 | na | 87.5 |
| No Drift | 81.6 | na | 83.6 |

In the Contaminants Purified case there isn’t much appreciable shift in mean plDDT across 20 interations of search. Likely the input Rep is part of a family with a strong evolutionary signal and any contamination never appreciably contributes to the AlphaFold2 performance. In the contaminants complex case having larger number of contaminating families does appear to reduce mean plDDT by about 7 points. This is likely a dense region of protein sequence space and families may have a wider degree of structural diversity as a consequence. Finally, when Contaminating families grow as a proportion of the results this appears to improve the results when the proportion of contamination families is at its peak. This may be due to the fact that the peak point of contamination is also the peak amount of sequence diversity in these cases. And In these cases there is typically only 1 or 2 off-target sequence families. Nevertheless the results show that AlphaFold2 is very robust to changes in the makeup of the MSA.

To further analyse the impact of the changing plDDT scores, merizo search was run for each of the Alphafold2 models and the predicted CATH structural domain ID was recorded. This data was then collated to (see table 5) to record how often the CATH Homlogous Superfamily (evolutionary family) stayed the same from the first to the final iteration of search and what the change in TM Score was. Table 5 summarises these results.

**Table 5:**
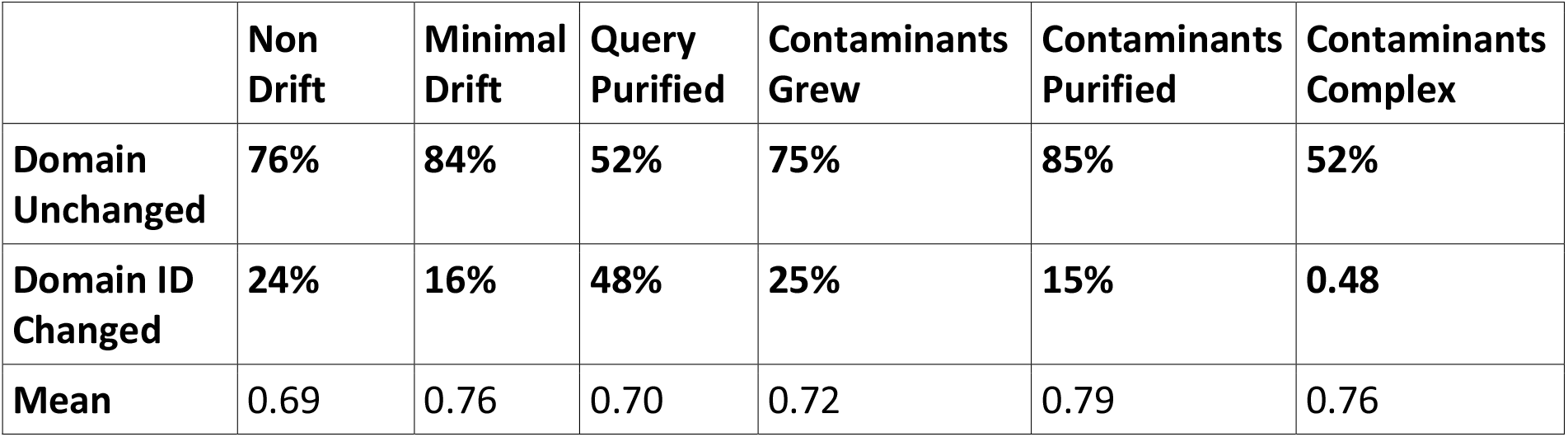
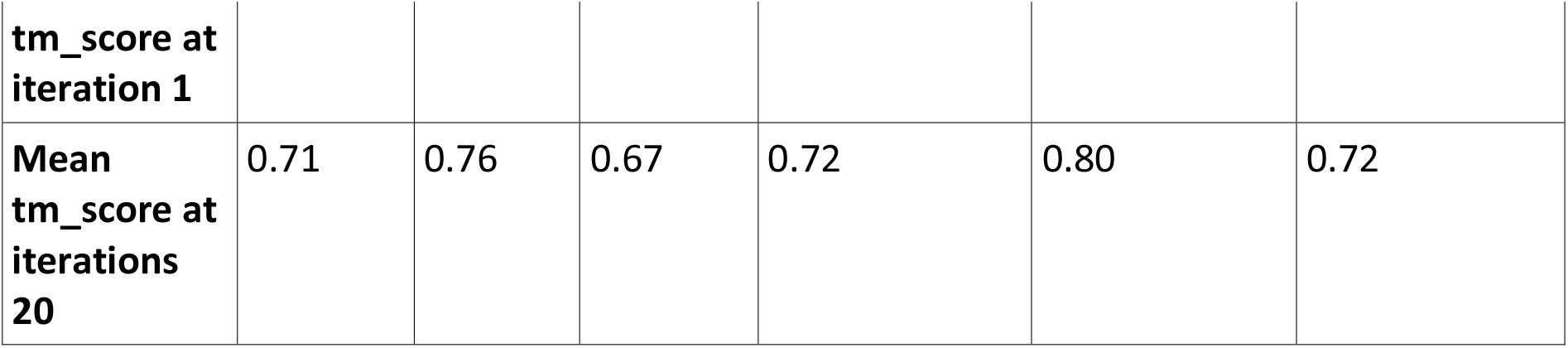
percentage of alphafold matches per drift class comparing the first and final iteration.

One important observation is that between 46 and 67% AlphaFold models built do not match a known CATH homologous super family (H-family). This is to be expected as the models were built using alignments of Pfam domain family members. Pfam domain and CATH H-families do not have one-to-one correspondence. Some sequence domains may represent more than one structural domain and some may represent just a portion of a structural domain. In Table 5 we present only results for the subset of analysed Pfam domains where there were matches to CATH families.

What can be seen from table 5 is that in the majority of cases the alphfold model remains matched to the same CATH-H family even as the make up of the MSA changes across the iterations. If we consider a structural domain family to be a “gold standard” indicator of evolutionary relatedness this may imply that many out-of-family matches are not out-of-family. They are instead wholly valid sequence matches in a dense region of sequence domain space and the fact the Pfam divides this in to multiple sequence families is likely an artefact of classification rather than a reflection of the sequence evolution. That is, these are not examples of the search drifting. Even in the non and minimal drift classes we see some shifts in H-family membership for the models. Perhaps this indicates that the CATH classification is dividing up structural space too finely. As such the “Domain Unchanged” row in table 5 may indicate that these examples of drift are more of a classification artefact than a statistical error in the search process.

Looking across the “Domain ID Changed” row these are instances where as the make up of the MSA is changing across each search iteration and changes to the structural models result in different CATH H family designations. These are likely the clearest examples of actual drift in the search. As expected, in the Non Drift, Minimal Drift and Contaminants purified cases this is least likely to happen as the MSA is typically dominated by sequence from the query sequence’s Pfam family. The query purified and contaminants complex classes are most likely to show structural change. In the first case there are no sequences from the query left at the end of search and in the second there are sequences from many differing families making the MSA composition complex. The Contaminants Grew class mirrors the behaviour of the non-drift class. In some cases there is possibly there is no drift here, and on average the Query family and Contaminant family are perhaps better considered part of the same evolutionary family. In other instances we see that the contaminant family is always the dominant family in the MSA in all iterations so the model likely reflects this.

### Coherence, hmm, and pLMs

A typical iterative search using a tool such as PSI-BLAST, HMMer or HHBlits is usually run for 5 or fewer iterations. At iteration 5 we calculated the percentage of sequence contaminants for each of the drift classes. This gives and way to directly measure how sequence family-specific the PSI-BLAST PSSM is at iteration 5. That is, how much of the mean variance at each column of the MSA is contributed to by in-family sequences and out-of-family sequences.

To explore how family-specific other sequence family representations may be this was compared to a Hidden Markov Model (HMM) and Protein Language Model (pLMs) representation. The PSI-BLAST hits were compared to generated sequences from the Pfam HMMs, ProtTransT5 sequences at 3 different levels of input masking. Table 6 shows the results of this comparison. The variance in these results should not be over-interpreted as the percentages are based only on an analysis of 100 Pfam families per drift class.

**Table 6:** Percentage of out of family hits per protein family representation.

| Drift class | PSI-BLAST | HMM | ProtTransT5 25% | ProtTransT5 50% | ProtTransT5 75% |
| --- | --- | --- | --- | --- | --- |
| Contaminants Grew | 63 | 19.5 | <0.1 | 0.6 | 37 |
| Contaminants Complex | 79 | 19.8 | <0.1 | 0.4 | 35 |
| Contaminants Purified | 8 | 19.5 | 0.1 | 0.7 | 28 |
| Query Purified | 88 | 17.9 | 0.2 | 0.7 | 40 |
| Minimal | <0.1 | 19.7 | <0.1 | 0.2 | 49 |
| No Drift | 0% | 18.3 | <0.1 | 0.2 | 49 |

Surveying these data the obvious standout is that regardless of drift category the HMM generated sequences do not belong to the HMMs sequence family in nearly 20% of cases. Here they either best hit a different family or do not align to a known sequence from a Pfam (approximately 8-10% of the time in either case). This is to be somewhat expected. A protein sequence HMM only attends to local relationships between columns in an MSA. Real sequences, due to physical effects such as protein folding(2), are constrained by long range contacts that can occur at any sequence separation and long range effects are not modelled by sequence HMMs. We would expect they would struggle to produce realistic in-family sequences as a consequence. It is notable how uniform this effect is regardless of drift category.

The ProtTransT5 classes clearly demonstrate the extent to which these models memorise their training data. When the masking percentage is below 50% the model almost always produces an in-family sequence. While developing this analysis, at 25% sequence masking, the model would frequently output the exact target input sequence indicating a fair degree of training data memorisation. This should be expected the masked sequence training task used to develop these pLLM embeddings is specifically designed to have the model memorise or hash the training data. It is interesting that the training model is very robust to large amounts of masking up to 50% well beyond the masking in the training task.

At high degrees of sequence masking, 75%, the model struggles more to produce in-family sequences often performing twice as poorly as the HMM model. This is interesting as analysis of database such as CATH indicate proteins in the same evolutionary family can have sequence identities as low as 9%. If the pLM had learnt the salient residues within each encoded sequence family should we expect better performance here? This results reinforces the notion that the network memorises the training input and does not necessarily learn the salient residues. And this should be expected as the loss function is not designed to penalise the network for failing to learn specific family-critical residues. Though this does suggest a potential improvement for protein family loss functions to weight the loss by the family relevant residues for each input sequence.

Overall the PSI-BLAST, HMM and highly masked ProtTransT5 protein family cases don’t indicate a high degree of protein family specificy encoded in the models, for this specific task. Though the moderate and low masked ProtTrans and minimal/no drift PSI-BLAST cases do indicate highly specificly family representations encoded in the family “model”.

## Conclusions Discussion

This paper investigated the occurrence of profile drift in iterative sequence search. Putative drift effects are seen in 15% of the searches performed (see Table 1) about a half to two thirds of these (Table 5) are likely artefacts of the Pfam family clustering and annotation process which leaves less than 5% of searches showing undesirable drift effects where search results are contaminated with out-of-family results and this can be backed up by changes in the predicted structures and structural families in these cases. For many analyses this degree of contamination will be acceptable. A single user making use of NCBI Blast online can easily appraise their results and use expert knowledge to discard unsuitable sequence hits.

For large scale projects which rely on automated high-throughput sequence search, such as AlphaFoldDB(3) or CATH, this may be an issue. Maximally, 1 in 20 contaminated MSAs may have impacts on downstream predictions and annotations. When performing many millions of sequence annotations this means a substantial number of annotations may be affected. However, this study looked only single protein domains. In realistic search cases many proteins have multiple domains and matching the full sequence will place additional constrains on the search that will constrain drift issues. Though for a project such as Interpro, Pfam and CATH, who *are* making assignments on single domains, this issue may remain significant.

One important observation is for the use of iterative search is that most changes in MSA composition are fixed by the 5^th^ iteration of search. This gives an empirical basis to the common practice of only running iterative search for fewer than 5 iterations.

Overall these observations should prove useful in the further development of better sequence search tools and for any analyses that rely on large scale sequence search. This includes compiling secondary bioinformatics databases (such as Interpro, Pfam and CATH) or using predictors such as AlphaFold2, PSIPRED(4) and MEGA(5)

